# Hyper-variability in Circulating Insulin Levels and Physiological Outcomes to High Fat Feeding in Male *Ins1*^−/−^:*Ins2*^+/−^ Mice in a Specific Pathogen-free Facility

**DOI:** 10.1101/031799

**Authors:** Nicole M. Templeman, Arya E. Mehran, James D. Johnson

## Abstract

Insulin is an essential hormone with key roles in energy homeostasis and body composition. Mice and rats, unlike other mammals, have two insulin genes: the rodent-specific *Ins1* gene and the ancestral *Ins2* gene. The relationships between insulin gene dosage and obesity has previously been explored in male and female *Ins2*^−/−^ mice with full or reduced *Ins1* dosage, as well as in female *Ins1*^−/−^ mice with full or partial *Ins2* dosage. We report herein unexpected hyper-variability in circulating insulin and physiological responses to high fat feeding in male *Ins1*^−/−^:*Ins2*^+/−^ mice. Two large cohorts of *Ins1*^−/−^*:Ins2*^+/−^ mice and their *Ins1*^−/−^:*Ins2*^+/+^ littermates were fed chow diet or high fat diet (HFD) from weaning and housed in specific pathogen-free (SPF) conditions. Cohort A and cohort B were studied one year apart. Contrary to female mice from the same litters, inactivating one *Ins2* allele on the complete *Ins1*-null background did not cause a consistent reduction of circulating insulin in male mice. In cohort A, HFD-fed males showed an equivalent degree of insulin hypersecretion and weight gain, regardless of *Ins2* dosage. In cohort B, *Ins1*^−/−^:*Ins2*^+/−^ males showed decreased insulin levels and body mass, compared to *Ins1*^−/−^:*Ins2*^+/+^ littermates. While experimental conditions were held consistent between cohorts, we found that HFD-fed *Ins1*^−/−^:*Ins2*^+/−^ mice with lower insulin levels had increased corticosterone. Collectively, these observations highlight the hyper-variability and range of phenotypic characteristics modulated by *Ins2* gene dosage, specifically in male mice.

## Introduction

Variations in circulating insulin levels have far-reaching metabolic consequences. In addition to the expected fluctuations in insulin secretion that are associated with blood glucose, circulating insulin levels are affected by a number of hormones and circulating factors, including amino acids, fatty acids, estrogen, melatonin, leptin, growth hormone, glucose-dependent insulinotropic polypeptide, and glucagon-like peptide-1 (see [1]). In mice, the mean 5-h fasted insulin levels in non-obese, 12 week-old males can range from 0.5 to 1.2 ng/mL, across four commonly used strains [2]. In humans, fasting insulin levels can range from 0.04 to 3.43 ng/mL in a nondiabetic adult population [3,4], and evidence suggests that less than half of the variance in fasting insulin can be can be accounted for by genetic variability [5,6].

Mice and rats have two non-allelic insulin genes, with a rodent-specific *Ins1* gene that likely arose from the transposition of a reverse-transcribed, partially processed mRNA of the ancestral *Ins2* [7]. *Ins1* and *Ins2* genes reside on different chromosomes in mice [8]. While *Ins1* lacks one of the two introns found in *Ins2,* the murine *Ins* genes share high homology up to 500 base pairs preceding the transcription initiation site [9]. *Ins1* and *Ins2* have distinct promoter elements, tissue- and temporal-specific expression patterns, and imprinting status [10–14]. In addition, differential translation or processing rates of the two murine preproinsulins have been reported [15,16]. It is therefore possible that levels of the fully processed murine insulin 1 and insulin 2 peptides are divergently susceptible to modulation under various conditions, which could underlie the evolutionary retention of both genes [17]. When one insulin gene is inactivated, elevated transcript and protein level of the non-deleted insulin gene can at least partially compensate for the loss [18], although the exact nature of this reciprocal relationship remains understudied.

We have performed a series of investigations to examine how murine *Ins1* and *Ins2* gene dosage impacts the onset of high fat diet-induced hyperinsulinemia and the development of obesity. Previous work in our laboratory showed that reducing *Ins1* gene dosage (on an *Ins2*-null background) results in continuous suppression of fasting hyperinsulinemia in male mice, thereby preventing diet-induced obesity [14]. Interestingly, circulating insulin levels were not similarly modulated in the female littermates from this study [14], suggesting the possibility of sex-specific differences in the relationship between insulin gene dosage and circulating insulin levels. In the converse genetic manipulation, reduced *Ins2* dosage (on an *Ins1*-null background) led to high-fat fed female *Ins1*^−/−^:*Ins2*^+/−^ mice having lower insulin secretion than their *Ins1*^−/−^:*Ins2*^+/+^ controls at a young age, which again corresponded with attenuated obesity [19]. The phenotype of the female *Ins1*^−/−^:*Ins2*^+/−^ mice was highly consistent between the two large cohorts of animals studied under specific pathogen-free (SPF) conditions, and was congruent to preliminary evaluations in a conventional facility.

We report herein on circulating insulin levels and the metabolic phenotype of the male *Ins1*^−/−^:*Ins2*^+/−^ and *Ins1*^−/−^:*Ins2*^+/+^ littermates of the female mice that were the subject of our recent investigation [19]. Contrary to our expectations, inactivating one *Ins2* allele did not cause a consistent reduction of circulating insulin in *Ins1*-null male mice, which precluded us from properly testing the hypothesis that reduced *Ins2* dosage and lower insulin levels would lead to protection from obesity in males. Specifically, we report that across cohorts, the effects of high fat feeding on glucose homeostasis, insulin sensitivity, and weight gain in *Ins1*^−/−^:*Ins2*^+/−^ male mice varied widely. Moreover, circulating insulin levels were hyper-variable across cohorts in *Ins1*-null male mice, pointing to sex-specific compensation of insulin homeostasis in these animals. Differences in degree of insulin compensation were associated with corticosterone levels, a marker of stress. Together with the accompanying study, we demonstrate that there is phenotypic hyper-variability within two different animal facilities (one conventional and one SPF). Collectively, these reports along with our previous published work [14,19] illustrate that the physiological outcomes of *Ins2* haploinsufficiency in male *Ins1*-null mice are dramatically variable relative to female littermates, and relative to mice with reduced *Ins1* gene dosage on an *Ins2*-null background. This phenotypic hyper-variability is evident despite the controlled environment of a SPF facility.

## Materials and Methods

### Experimental Animals, Body Composition and Metabolic Cage Experiments

All animal procedures were approved by the University of British Columbia Animal Care Committee, following Canadian Council for Animal Care guidelines. Mice with *Ins1*-null and *Ins2*-null alleles were generated previously [20] and were roughly equal parts C57BL/6 and 129 background. Data presented here are from the male littermates of previously described *Ins1*^−/−^:*Ins2*^+/+^ and *Ins1*^−/−^:*Ins2*^+/−^ females [19], and were therefore collected in the same time-frames and conditions. All animals were predominately handled by the same female researcher. The mice tracked across time were in two major cohorts, born a year apart (cohort A, born October 2011 – December 2011, and cohort B, born October 2012 – February 2013; Fig. 1a). The dams and sires of cohort B experimental mice were themselves born of parents from cohort A stock, thus minimizing the chance of significant genetic drift, and the same room was used for breeding across both cohorts. Shortly after weaning, mice were assigned a moderate-energy chow diet (CD, 25% fat content; LabDiet 5LJ5; PMI Nutrition International, St. Louis, MO, USA) or a high-energy high fat diet (HFD, 58% fat content; Research Diets D12330; Research Diets, New Brunswick, NJ, USA) provided *ad libitum,* except during fasting periods (Fig. 1a). Diet assignments were distributed within each litter, based on approximately matching starting body weights between diet groups. Mice were housed under SPF conditions at 21°C, on a 12:12 h light:dark cycle, in the same room for both cohorts. The vast majority of male mice from both cohorts were individually housed, due to fighting between young cage-mates.

**Figure 1.**
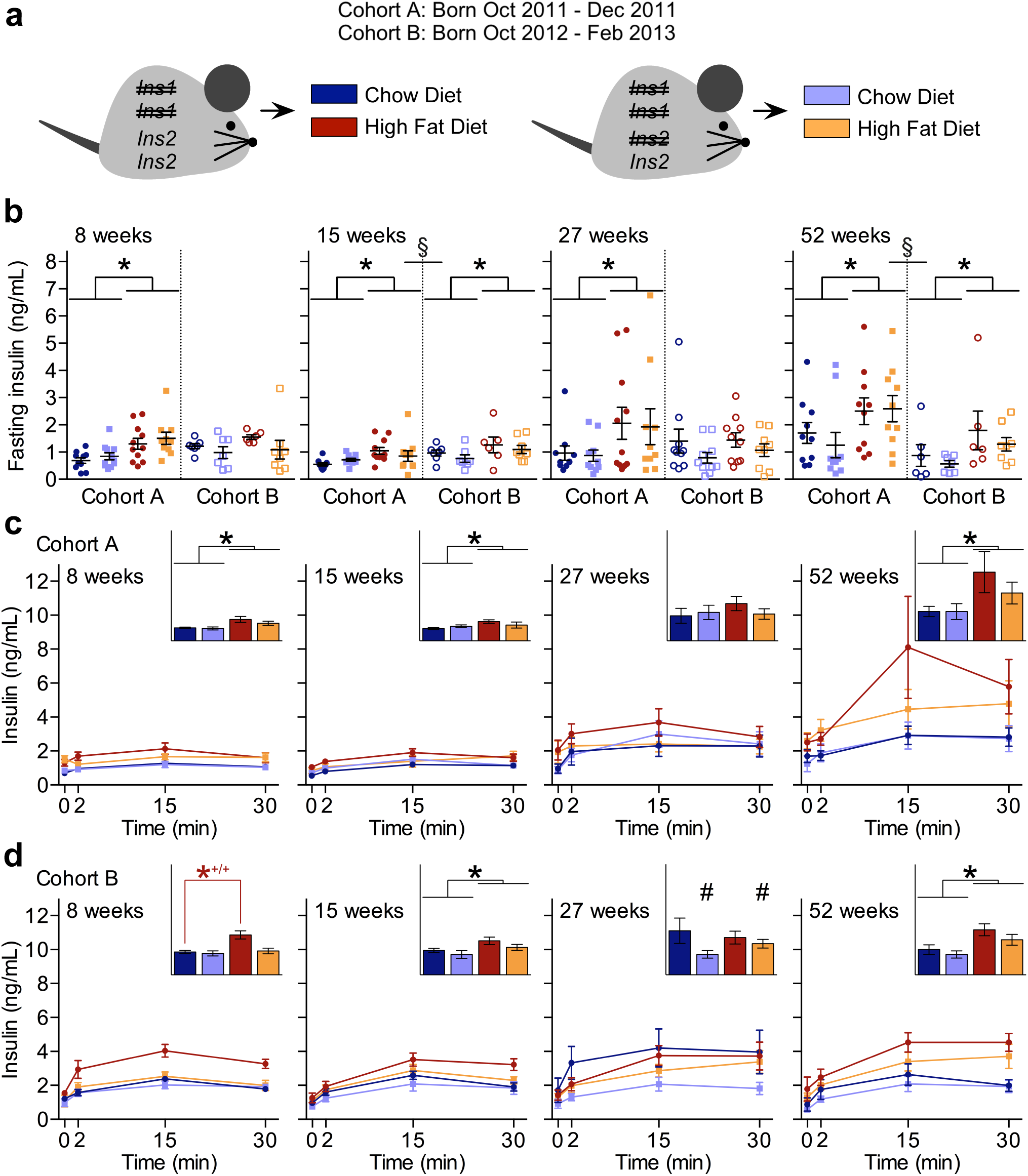
Hyper-variability in Insulin Secretion Across Two Cohorts. (**a**) Experimental design showing two cohorts (A and B) of *Ins1*^−/−^:*Ins2*^+/+^ and *Ins1*^−/−^:*Ins2*^+/−^ male littermates fed chow diet (CD) or high fat diet (HFD). (**b**) Periodic measurements of 4-h fasted insulin levels is shown for both cohorts, with scatter points to indicate individual values (cohort A: closed points, n = 10−11; cohort B: open points, n = 6−10). In addition, periodic measurements of glucose-stimulated insulin secretion in (**c**) cohort A (n = 10−11) and (**d**) cohort B (n = 6−8) is shown, with area under the curve (y-axis units of ng/mL•min) in panel insets. Data are means ± SEM. Dark blue and red represent CD- and HFD-fed *Ins1*^−/−^:*Ins2*^+/+^ male mice, respectively; pale blue and orange represent CD- and HFD-fed *Ins1*^−/−^:*Ins2*^+/−^ male mice, respectively. *p* ≤ 0.05 denoted by * for CD vs HFD, *^+/+^ for CD-vs HFD-fed *Ins1*^−/−^:*Ins2*^+/+^ mice, # for *Ins1*^−/−^:*Ins2*^+/+^ vs *Ins1*^−/−^:*Ins2*^+/−^ § for cohort A vs cohort B.

After weaning, all mice were weighed weekly, although a subset of pups from cohort B was weighed earlier. *In vivo* body composition was periodically assessed in cohort B mice using dual-energy X-ray absorptiometry (DEXA; Lunar PIXImus densitometer; GE Medical Systems LUNAR, Madison, WI, USA). A subset of 17 week-old HFD-fed mice from cohort B was evaluated in PhenoMaster metabolic cages (TSE Systems, Chesterfield, MO, USA) as described [21] after individual housing for at least a week, and acclimation to the metabolic cage environment for at least four days. Values were averaged from 48–84 h of continual data acquisition.

### Glucose Homeostasis and Plasma Analytes

Mice were fasted for 4 h during the light period to ensure a postprandial state for all blood sampling. Fasting and glucose-stimulated (2 g/kg) insulin secretion was assessed, as well as blood glucose response to intraperitoneal delivery of glucose (2 g/kg) or an insulin analog (0.75 U/kg of Humalog; Eli Lilly, Indianapolis, IN, USA). OneTouch Ultra2 glucose meters (LifeScan Canada Ltd, Burnaby, BC, Canada) were used for blood glucose assessments, and plasma insulin was measured with a mouse insulin ELISA (Alpco Diagnostics, Salem, NH, USA), according the manufacturers’ instructions. Area over or under the curve was calculated to evaluate statistical differences in these glucose- or insulin-stimulated tests. Corticosterone was measured in plasma collected in the early afternoons across multiple weeks for each cohort, and assessed using a mouse/rat corticosterone ELISA (Alpco Diagnostics) according the manufacturers’ instructions. In plasma from 40 week-old cohort B mice, we used colorimetric kits to measure total cholesterol (Cholesterol E kit; Wako Chemicals, Richmond, VA, USA), triglycerides (Serum Triglyceride kit; Sigma-Aldrich, St Louis, MO, USA), and non-esterified fatty acid levels (NEFA-HR[2] kit; Wako Chemicals), in addition to a mouse magnetic bead panel assay utilizing Luminex technology (Millipore, St. Charles, MO, USA) to evaluate leptin, resistin, interleukin 6, glucose-dependent insulinotropic polypeptide, peptide YY, and glucagon levels, in accordance with manufacturers’ instructions.

### Islet Analyses

Islet *Ins2* mRNA, insulin content, and function were assessed in mice that were discrete from the other experimental groups, as the A and B cohorts were assessed longitudinally. 25 week-old mice (born April 2014) were euthanized after 4 h of fasting in the light period, and HFD-fed 70 week-old mice (born April 2012) were euthanized after a minimum of 1 h of fasting. Islets were isolated with collagenase and filtration, using a previously described protocol [22] with minor modifications. Islets were handpicked using brightfield microscopy, and cultured at 37°C and 5% CO_2_ for at least 16 h prior to any analyses, in RPMI-1640 medium (Invitrogen, Burlington, ON, Canada) supplemented with 11 mM glucose, 100 U/mL penicillin, 100 μg/mL streptomycin and 10% (volume/volume) fetal bovine serum. Islet perifusion experiments were carried out as previously described [23], using groups of 150 size-matched islets and evaluating stimulatory conditions of 15 mM glucose or 30 mM KCl. Islet insulin content were averaged from 10 size-matched islets per mouse, incubated at −20°C in 0.02 M HCl in 70% ethanol, and sonicated a minimum of 30 s before dilution for measurement with a mouse insulin ELISA kit (Alpco Diagnostics).

Quantitative reverse transcription PCR was used for the relative quantification of *Ins2* in islets, normalized against the reference gene ß-actin. Total RNA was extracted from islets using the Qiagen RNeasy Mini kit (Qiagen, Mississauga, ON, Canada), according to manufacturers’ instructions. RNA was quantified at 260 nm with a NanoDrop 2000 spectrophotometer (Thermo Scientific, Wilmington, DE, USA), and cDNA was generated using a qScript cDNA synthesis kit (Quanta Biosciences, Gaithersburg, MD, USA). Transcript levels were measured with Taqman assays (*Ins2*: forward primer 5'-GAAGTGGAGGACCCACAAGTG-3', reverse primer 5'-GATCTACAATGCCACGCTTCTG-3'; Integrated DNA Technologies, Toronto, ON, Canada; ß-actin: assay catalog number 4352341E, Applied Biosystems, Foster City, CA, USA). Reactions were performed in duplicate on a StepOnePlus real-time PCR system (Applied Biosystems), using TaqMan Fast Universal PCR Master Mix (Applied Biosystems) and a fast-mode thermal program consisting of a 20-s activation at 95°C, then 40 cycles of 95°C melting for 1 s and 60°C annealing for 20 s. Values were normalized using the 2^−Δ^*^Ct^* method.

### Statistical Analyses

Statistical analyses were performed with SPSS 15.0 software, and a critical α-level of *p* ≤ 0.05. For most analyses, we used two-way analysis of variance (ANOVA) models within each cohort to assess factors of genotype and diet, and a significant interaction led to one-way ANOVAs comparing HFD-fed *Ins1*^−/−^:*Ins2*^+/+^ mice, CD-fed *Ins1*^−/−^:*Ins2*^+/+^ mice, HFD-fed *Ins1*^−/−^:*Ins2*^+/−^ mice, and CD-fed *Ins1*^−/−^:*Ins2*^+/−^ mice, with Bonferroni corrections. Three-way ANOVAs were used for incorporating the additional factor of cohort or parental effect, and 2-tailed independent *t*-tests were used to assess differences if there were only two groups to be compared. Analysis of covariance was employed to test energy expenditure with covariates of lean and fat mass. In all cases, we used Levene’s test to evaluate the assumption of homogeneity of variance, and where the assumption was violated, logarithmic transformations were applied, generally stabilizing data variance. Graphpad Prism 6.0 software was used to generate and assess the linear regressions.

## Results

### Variability in Insulin Secretion Between Cohorts of Male Mice

We have recently studied the effects of reducing *Ins2* gene dosage in *Ins1*-null female mice, and found consistent outcomes for insulin secretion and body composition across cohorts [19]. While characterizing male siblings from the same cohorts, we noticed that a number of measured parameters showed dramatic inconsistencies between cohorts A and B, precluding us from pooling the data from these two cohorts. For instance, we observed that in cohort A males, fasting insulin was significantly higher in HFD-fed mice than CD-fed mice at all measured time points across a year (Fig. 1b), and glucose-stimulated insulin secretion was higher at 8, 15, and 52 weeks of age (Fig. 1c). Unexpectedly, there were no significant differences in circulating insulin levels (either fasting or glucose-stimulated) between *Ins1*^−/−^:*Ins2*^+/+^ and *Ins1*^−/−^:*Ins2*^+/−^ mice, at any point up to one year in cohort A (Figs. 1b,c). In contrast, cohort B *Ins1*^−/−^:*Ins2*^+/−^ mice tended to have lower fasting insulin levels than their *Ins1*^−/−^:*Ins2*^+/+^ littermates (Fig. 1b), and at 27 weeks of age they had significantly reduced glucose-stimulated insulin secretion (Fig. 1d). In addition, *Ins1*^−/−^:*Ins2*^+/+^ mice were the only cohort B males showing more glucose-stimulated insulin secretion on HFD than CD at the early 8-week time point (Fig. 1d). However, the HFD-induced elevation of basal and glucose-stimulated insulin secretion was, in general, quite modest for most males in cohort B (Figs. 1b,d). Notably, by 52 weeks all groups of cohort B male mice clearly had lower average insulin levels than cohort A males (Fig. 1b).

To evaluate potential mechanisms underlying cross-cohort changes in circulating insulin levels in male mice, we assessed insulin mRNA, protein, and islet function in two distinct groups. Due to the longitudinal nature of the experiments, we could not use islets from cohort A or B mice. However, a separate group of 25 week-old male mice tended to show a genotype effect for *in vivo* fasting insulin (*p* = 0.056), as well as similar raw insulin values to those of cohort B mice (Figs. 1b, 2a). Islets from these 25 week-old *Ins1*^−/−^:*Ins2*^+/−^ mice had an expected reduction in *Ins2* mRNA compared to *Ins1*^−/−^:*Ins2*^+/+^ islets (Figs. 2b). Interestingly, the significant reduction in *Ins2* mRNA did not correspond to significant genotype differences in islet insulin protein content (Fig. 2c), suggesting the involvement of post-transcriptional compensation. Consistent with the lack of a difference between genotypes for islet insulin content, dynamic secretion by islets from 25 week-old HFD-fed *Ins1*^−/−^:*Ins2*^+/−^ mice was not reduced compared to *Ins1*^−/−^:*Ins2*^+/+^ islets, and in fact 25 week-old *Ins1*^−/−^:*Ins2*^+/−^ islets appeared to have the capacity for a marginally increased 2^nd^ phase response to KCl stimulation (Fig. 2d).

**Figure 2.**
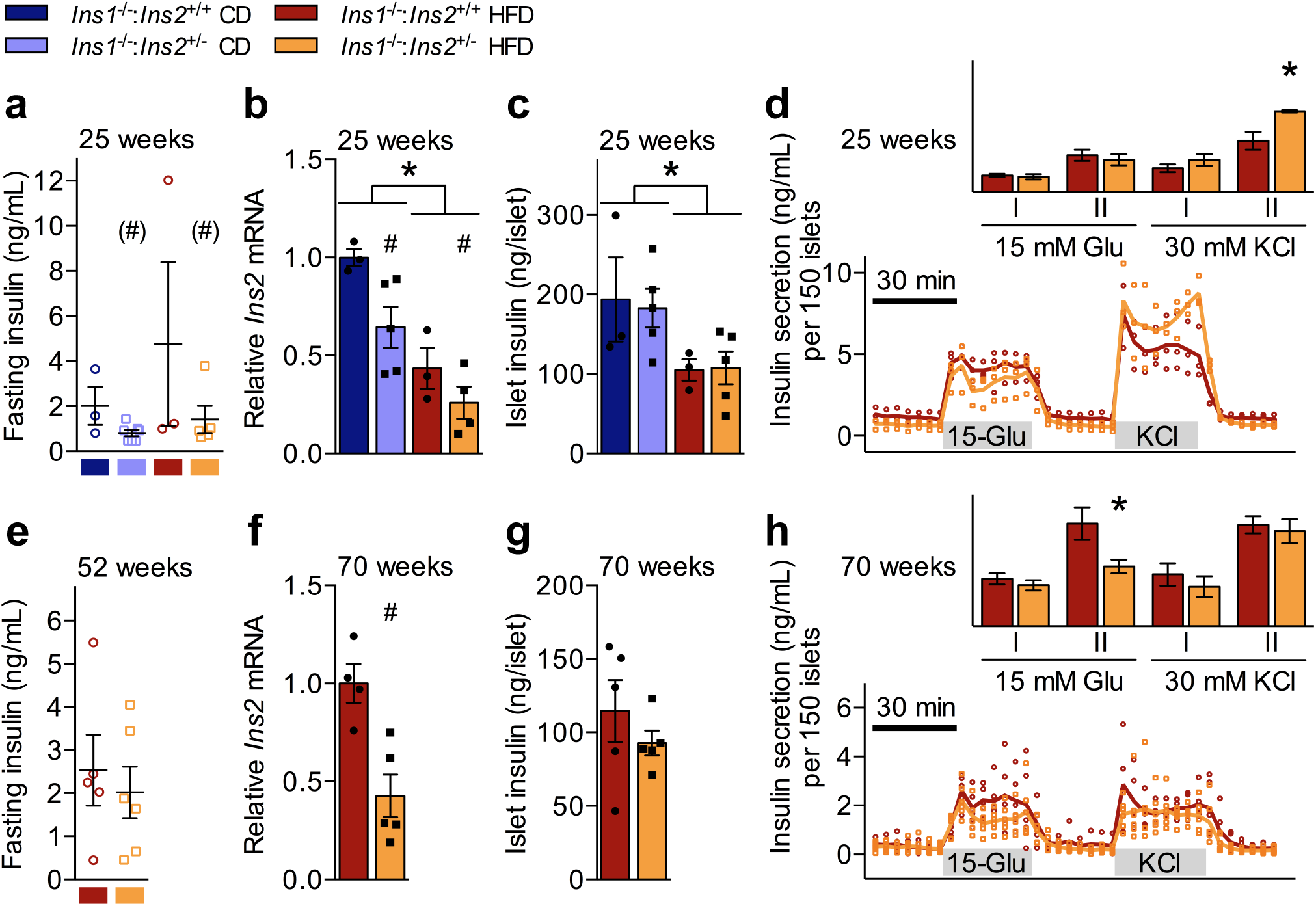
Characterization of *Ins1*^−/−^:*Ins2*^+/+^ and *Ins1*^−/−^:*Ins2*^+/−^ Islets. (**a**) *In vivo* 4-h fasted insulin is shown from 25 week-old mice prior to collection of islets to assess (**b**) *Ins2* mRNA, corrected against *ß-actin* and normalized to CD-fed *Ins1*^−/−^:*Ins2*^+/+^ mice, (**c**) islet insulin content, and (**d**) insulin secretion by 150 islets perifused with 3 mM glucose (basal), 15 mM glucose (Glu), and 30 mM KCl, with area under the curve (y-axis units of μg/mL*min) depicted for phases I/II of glucose and KCl stimulation, minus basal secretion. (**e**) *in vivo* 4-h fasted insulin is shown from 52 week-old mice prior to collection of islets at 70 weeks to assess (**f**) *Ins2* mRNA, corrected against *ß-actin* and normalized to HFD-fed *Ins1*^−/−^:*Ins2*^+/+^ mice, (**g**) islet insulin content, and (**h**) insulin secretion by 150 islets perifused with 3 mM glucose (basal), 15 mM glucose (Glu), and 30 mM KCl, with area under the curve (y-axis units of ng/mL•min) depicted for phases I/II of glucose and KCl stimulation, minus basal secretion. n = 3−7. Data are means ± SEM, with scatter points to indicate individual values. Dark blue and red represent CD- and HFD-fed *Ins1*^−/−^:*Ins2*^+/+^ male mice, respectively; pale blue and orange represent CD- and HFD-fed *Ins1*^−/−^:*Ins2*^+/−^ male mice, respectively. *p* ≤ 0.05 denoted by * for CD vs HFD, # for *Ins1*^−/−^:*Ins2*^+/+^ vs *Ins1*^−/−^:*Ins2*^+/−^ (#) indicates *p* = 0.056 for *Ins1*^−/−^:*Ins2*^+/−^ vs *Ins1*^−/−^:*Ins2*^+/−^.

We also evaluated 70 week-old islets from a group of HFD-fed mice that had shown no obvious genotype differences in fasting insulin levels at 52 weeks of age (Fig. 2e). This allowed us to confirm that the genetic manipulation also led to reduced *Ins2* mRNA in older HFD-fed *Ins1*^−/−^:*Ins2*^+/−^ male mice (Fig. 2f), with a similar capacity for compensation at the level of islet insulin content (Fig. 2g) as was evident at 25 weeks (Fig. 2c). The only detected differences in dynamic *in vitro* islet secretion were minimal, showing that at 70 weeks, *Ins1*^−/−^:*Ins2*^+/−^ islets did not secrete quite as much insulin in the 2^nd^ phase of glucose stimulation as *Ins1*^−/−^:*Ins2*^+/+^ islets (Fig. 2h). Collectively, these data show that although the experimental genetic manipulation did successfully reduce *Ins2* expression in the male mice, there was evidence of compensation with respect to insulin protein content and islet secretory capacity. The capability for insulin production and/or secretory compensation may have accounted for the lack of consistent differences in circulating insulin levels between *Ins1*^−/−^:*Ins2*^+/+^ and *Ins1*^−/−^:*Ins2*^+/−^ male mice.

### Variability in Glucose Homeostasis Between Cohorts of Male Mice

We also observed heterogeneity between cohorts in a longitudinal analysis of glucose homeostasis, as might be expected from cross-cohort variability in insulin levels. In conjunction with the sustained HFD-induced elevation of circulating insulin in cohort A mice, from 14 weeks onwards HFD-fed males from cohort A showed significant whole-body insulin resistance compared to their CD-fed littermates (Fig. 3a). HFD-fed mice in cohort A also had a modest degree of glucose intolerance compared to mice on CD, at 8 and 50 weeks (Fig. 3b). In contrast, there were no statistically significant differences in insulin sensitivity or glucose tolerance observed between CD- and HFD-fed groups of cohort B mice, consistent with a limited response to HFD-feeding (Figs. 3c,d).

**Figure 3.**
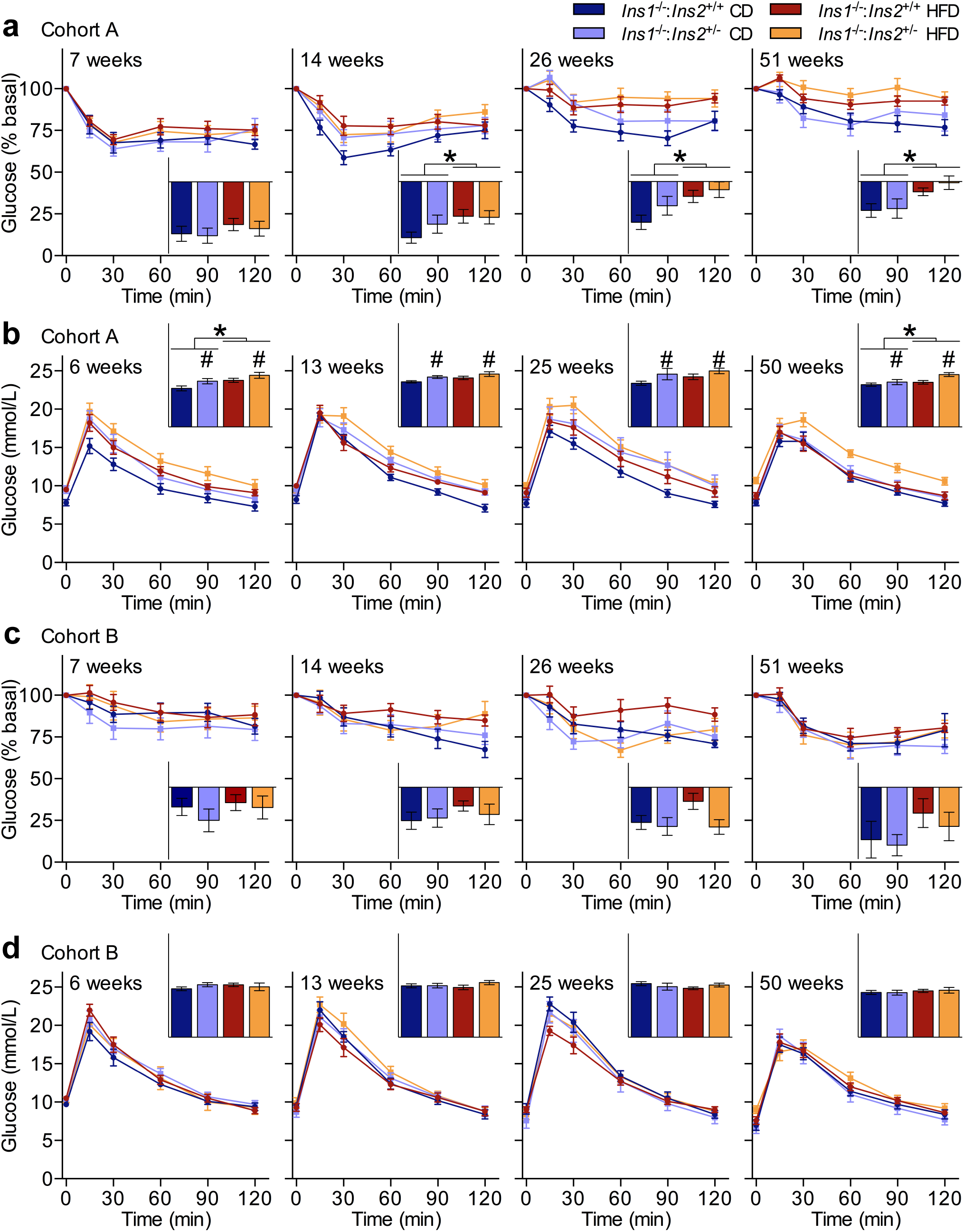
Variability in Glucose Homeostasis Across Two Cohorts. Periodic measurements of blood glucose response to intraperitoneal delivery of (**a**) an insulin analog (n = 10−19) in cohort A mice, and (**b**) glucose (n = 11−19) in cohort A mice, as well as (**c**) an insulin analog (n = 7−11) in cohort B mice, and (**d**) glucose (n = 7−11) in cohort B mice. Area under or over the curve (y-axis units of **a**,**c** percent•min, **b**,**d** mmol/L•min) is shown in panel insets. Data are means ± SEM. Dark blue and red represent CD- and HFD-fed *Ins1*^−/−^:*Ins2*^+/+^ male mice, respectively; pale blue and orange represent CD- and HFD-fed *Ins1*^−/−^:*Ins2*^+/−^ male mice, respectively. *p* ≤ 0.05 denoted by * for CD vs HFD, # for *Ins1*^−/−^:*Ins2*^+/+^ vs *Ins1*^−/−^:*Ins2*^+/−^.

Reduced *Ins2* dosage and the slight differences in circulating insulin levels observed between *Ins1*^−/−^:*Ins2*^+/+^ and *Ins1*^−/−^:*Ins2*^+/−^ cohort B mice did not appear to cause robust negative repercussions for glucose homeostasis (Figs. 3c,d). However, in cohort A, *Ins1*^−/−^:*Ins2*^+/−^ mice were slightly but significantly more glucose intolerant than their *Ins1*^−/−^:*Ins2*^+/+^ littermate controls at each measured time point across a year. Closer examination of the responses to intraperitoneal glucose stimulation shows a trend for a delayed or sustained peak in blood glucose in cohort A *Ins1*^−/−^:*Ins2*^+/−^ male mice (Fig. 3b). This suggests that although cumulative insulin levels appeared nearly matched between genotypes in cohort A, at least across the first 30 minutes post-stimulation (Figs. 1b,c), subtle differences in insulin secretory patterns (*e.g.* Fig. 2h, although this *in vitro* insulin secretion was assessed in different mice) could have affected glycemic control. Indeed, in addition to the long-term elevation of fasting glucose evident in HFD-fed mice compared to CD-fed mice in cohort A, there were periods where cohort A *Ins1*^−/−^:*Ins2*^+/−^ males had higher fasting glucose levels than their *Ins1*^−/−^:*Ins2*^+/+^ littermates (Fig. 4a). On the other hand, all groups of cohort B mice had largely comparable fasting glucose levels throughout the year (Fig. 4b).

**Figure 4.**
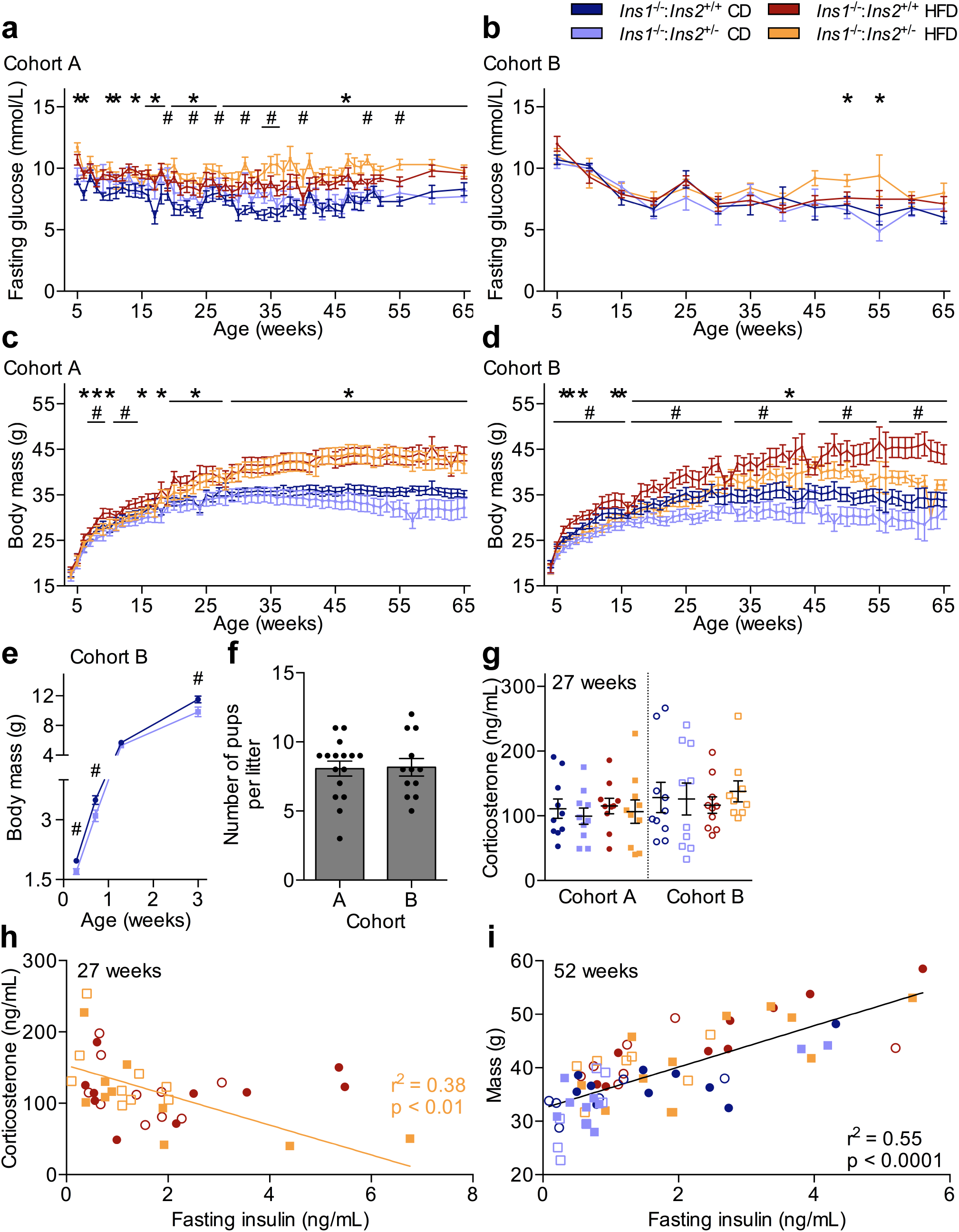
Cross-cohort Variation in Fasting Glucose, Body Mass, and Corticosterone. Periodic measurements of 4-h fasted blood glucose in (**a**) cohort A (n = 12−18, most time points) and (**b**) cohort B (n = 8−10, most time points) is shown, in addition to weekly body mass in (**c**) cohort A (n = 12−18, most time points) and (**d**) cohort B (n = 8−10, most time points). (**e**) Body mass was also tracked in a subset of pre-weaned male pups from cohort B (n = 12−22). (**f**) Litter sizes were compared between cohorts, with scatter points indicating the number of pups per individual litter (n = 12−16). 4-h fasted corticosterone levels, measured in plasma collected during early afternoon from 27 week-old mice, were assessed (**g**) between cohorts and (**h**) across cohorts, comparing HFD-fed mice with low fasted insulin (< 1.90 ng/mL) to HFD-fed mice with high fasted insulin (≥ 1.90 ng/mL), with scatter points to indicate individual values (cohort A: closed points, n = 10; cohort B: open points, n = 9−10). Data are means ± SEM. In addition, (**i**) The relationship between body mass and 4-h fasted insulin levels at one year of age is shown, with r^2^ = 0.55 and *p* < 0.0001 (cohort A: closed points, n = 9−12; cohort B: open points, n = 6−7). Dark blue and red represent CD- and HFD-fed *Ins1*^−/−^:*Ins2*^+/+^ male mice, respectively; pale blue and orange represent CD- and HFD-fed *Ins1*^−/−^:*Ins2*^+/−^ male mice, respectively. *p* ≤ 0.05 denoted by * for CD vs HFD, # for *Ins1*^−/−^:*Ins2*^+/+^ vs *Ins1*^−/−^:*Ins2*^+/−^, †^+/−^ for high vs low insulin *Ins1*^−/−^:*Ins2*^+/−^ groups.

### Heterogeneity Between Cohorts in Body Mass of *Ins1*^−/−^:*Ins2*^+/−^ Male Mice

Perhaps one of the most striking differences noted between cohorts A and B was the differential impact of reducing *Ins2* gene dosage on body weights, particularly in HFD-fed mice. The loss of one *Ins2* allele did not notably affect either circulating insulin levels (Figs. 1b,c) or HFD-induced growth (Fig. 4c) in cohort A males, since all HFD-fed mice showed equivalent weight gain compared to CD-fed littermates, particularly from 20 weeks onward (Fig. 4c). Although young *Ins1*^−/−^:*Ins2*^+/−^ mice were smaller than their *Ins1*^−/−^:*Ins2*^+/+^ littermates for a limited period in cohort A, this did not persist (Fig. 4c). HFD-fed mice were heavier than CD-fed mice for a similar duration in cohort B as in cohort A, but the significantly smaller mass of cohort B *Ins1*^−/−^:*Ins2*^+/−^ mice compared to their *Ins1*^−/−^:*Ins2*^+/+^ littermates continued throughout most of the year (Fig. 4d). Notably, a difference in body mass between *Ins1*^−/−^:*Ins2*^+/+^ and *Ins1*^−/−^:*Ins2*^+/−^ mice of cohort B was detectable in male pups as young as 2 days of age (Fig. 4e), and it is possible that there could have also been similar size differences in neonatal pups of cohort A (not measured) that might have contributed to the genotype effect on body mass observed in young cohort A mice (Fig. 4c).

We attempted to evaluate different factors that could have contributed to phenotypic differences between cohorts A and B. We did not detect obvious means by which parental imprinting of the *Ins2* allele might have accounted for the observed variability, as both breeding pair options (*i.e.* either the dam or the sire donating the disrupted *Ins2* allele) contributed to cohort A and B experimental animals. There were no statistically significant differences between offspring of dams versus sires with the disrupted *Ins2* allele when the factor of “parental effect” was incorporated into analyses of body mass or fasting insulin levels (with both cohorts pooled); any patterns suggestive of a possible parental effect in one cohort were either not present or showed opposite trends in the other cohort (data not shown). There was also no apparent distinction in the mean or distribution of litter sizes between cohorts A and B (Fig. 4f).

As a limited indicator of environmental stressors, we also measured 4 h-fasted corticosterone levels at 27 weeks. Interestingly, there tended to be a greater number of mice with elevated corticosterone levels in cohort B, and modest trends suggested that there were higher average corticosterone levels in HFD-fed *Ins1*^−/−^:*Ins2*^+/−^ males from cohort B, compared to cohort A (Fig. 4g). Closer examination of HFD-fed animals showed that across both cohorts, fasting insulin levels in HFD-fed *Ins1*^−/−^:*Ins2*^+/−^ male mice were inversely correlated to corticosterone levels (r^2^ = 0.38, *p* < 0.01), and this relationship was not evident in HFD-fed *Ins1*^−/−^:*Ins2*^+/+^ mice (r^2^ = 0.00, *p* = 0.95; Fig. 4h). Interpretation of these data is limited, as it is based on using a single measurement of corticosterone for each individual animal as a marker of ‘stress’ during an extended time period. However, it appears that those HFD-fed *Ins1*^−/−^:*Ins2*^+/−^ male mice that reached the highest fasting insulin levels at 27 weeks (predominately individuals from cohort A) tended to have lower corticosterone levels, whereas increased stress may have dampened the capacity of HFD-fed *Ins1*^−/−^:*Ins2*^+/−^ male mice to compensate for reduced *Ins2* dosage at the level of basal insulin secretion. If mice in cohort B did experience more stressful conditions than cohort A animals (Fig. 4g), this could be one of potentially many contributing factors underlying phenotypic differences in circulating insulin levels between cohorts A and B.

Heterogeneity in our data precluded pooling cohorts A and B to generate averaged results. However, although experimental genetic and dietary manipulations did not have consistent effects in both cohorts, we did observe a comparable range of fasting insulin values and body masses for both cohorts (Figs. 1b, 4c,d). To indirectly evaluate whether differences in fasting insulin might be underlying body weight alterations in this model, we examined the relationship between these two variables across all year-old mice. There was a positive correlation between body mass and fasting insulin levels at this age (r^2^=0.55, *p*<0.0001; Fig. 4i). Therefore, while we did not observe consistent effects of reducing *Ins2* gene dosage on circulating insulin and obesity in male mice, in general, these data support the concept that reduced insulin levels are associated with attenuated body weight and obesity.

### Attenuated Obesity and Fat-free Mass in Cohort B *Ins1*^−/−^:*Ins2*^+/−^ Males

We further characterized cohort B mice, as they showed a sustained divergence in body mass between *Ins1*^−/−^:*Ins2*^+/−^ and *Ins1*^−/−^:*Ins2*^+/+^ groups (Fig. 4d). First, we used metabolic cages to examine the *in vivo* energy balance of HFD-fed mice at 17 weeks, an age when both cohorts showed similar trends for body masses. Although HFD-fed *Ins1*^−/−^:*Ins2*^+/−^ mice exhibited a slight elevation in activity levels during the early hours of the dark period (Fig. 5a), it did not appear that there were genotype differences in whole-body energy expenditure (Fig. 5b), respiratory exchange ratio (Fig. 5c), or food intake (Fig. 5d) to account for the disparities in weight gain between HFD-fed *Ins1*^−/−^:*Ins2*^+/−^ and *Ins1*^−/−^:*Ins2*^+/+^ males.

**Figure 5.**
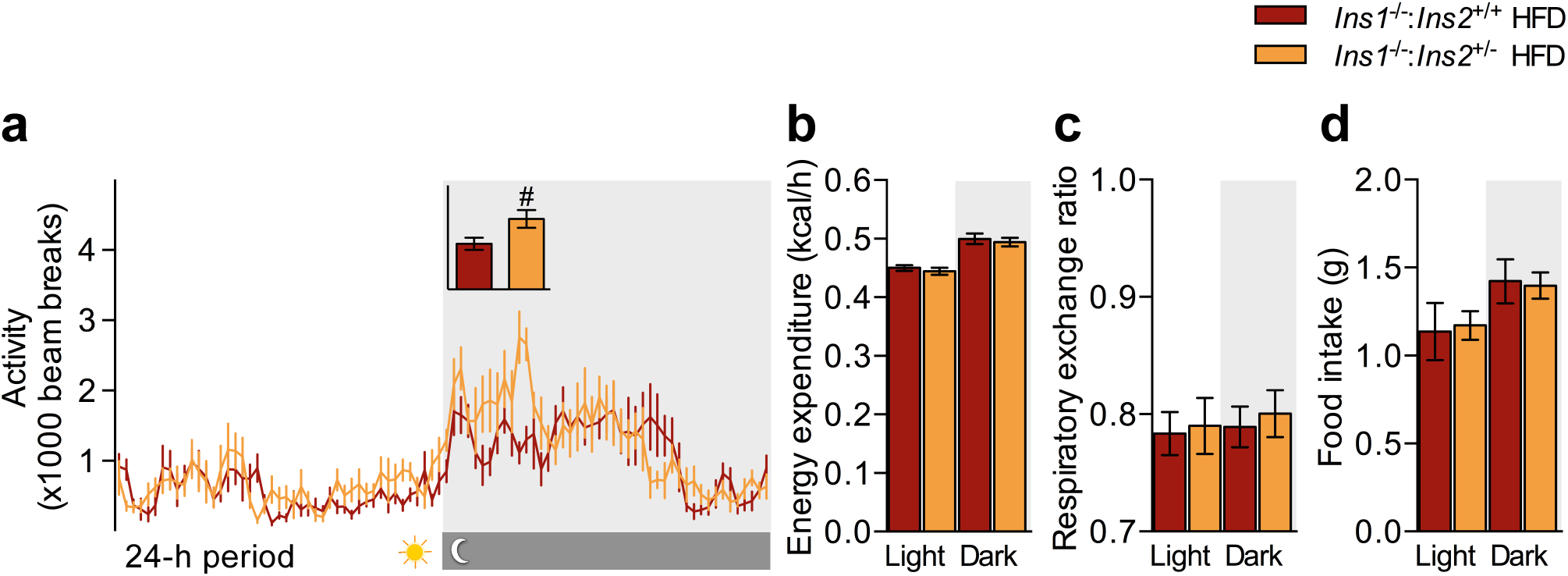
*In Vivo* Energy Homeostasis of HFD-fed Cohort B Males. In HFD-fed 17 week-old mice (n = 6−7), (**a**) 24-h activity, with inset showing mean beam breaks across the first 4 h of the dark period (y-axis units of beam breaks), as well as (**b**) energy expenditure, (**c**) respiratory exchange ratio, and (**d**) food intake were averaged across 48–84 h, with grey indicating dark cycles. Energy expenditure is shown as estimated marginal means ± SEM, adjusted for covariates of lean and fat mass; other data are simple means ± SEM. Dark red represents HFD-fed *Ins1*^−/−^:*Ins2*^+/+^ male mice; orange represents HFD-fed *Ins1*^−/−^:*Ins2*^+/−^ male mice. *p* ≤ 0.05 denoted by # for *Ins1*^−/−^:*Ins2*^+/+^ vs *Ins1*^−/−^:*Ins2*^+/−^.

We assessed body composition longitudinally with DEXA. Consistent with the evidence suggesting that lower *Ins2* dosage led to generally reduced growth in male neonatal pups (Fig. 4e), reductions in both adiposity and fat-free mass contributed to the smaller size of cohort B *Ins1*^−/−^:*Ins2*^+/−^ male mice. *Ins1*^−/−^:*Ins2*^+/+^ animals had significantly higher fat mass than *Ins1*^−/−^:*Ins2*^+/−^ males on both diets, at all measured time points (Fig. 6a). A similar pattern was also evident for fat-free masses, particularly at the older ages (Fig. 6b). However, since reductions in fat mass were proportionally greater than reductions in fat-free mass for *Ins1*^−/−^:*Ins2*^+/−^ versus *Ins1*^−/−^:*Ins2*^+/+^ males up to 60 weeks of age (Figs. 6a,b), it is clear that an attenuation in adiposity contributed to the reduced body mass of cohort B *Ins1*^−/−^:*Ins2*^+/−^ male mice, compared to their *Ins1*^−/−^:*Ins2*^+/+^ controls. In the group of 25 week-old males with similar insulin levels as cohort B mice (Figs. 1b, 2a), both subcutaneous and visceral white adipose tissue depots were smaller in *Ins1*^−/−^:*Ins2*^+/−^ versus *Ins1*^−/−^:*Ins2*^+/+^ mice, in addition to the reduced sizes of these depots in CD-mice compared to HFD-fed mice (Fig. 6c). Furthermore, the *Ins1*^−/−^:*Ins2*^+/−^ mice had smaller interscapular brown adipose tissue depots than their *Ins1*^−/−^:*Ins2*^+/+^ littermates, and a slight reduction in the size of a mixed triceps surae muscle group (Fig. 6c).

**Figure 6.**
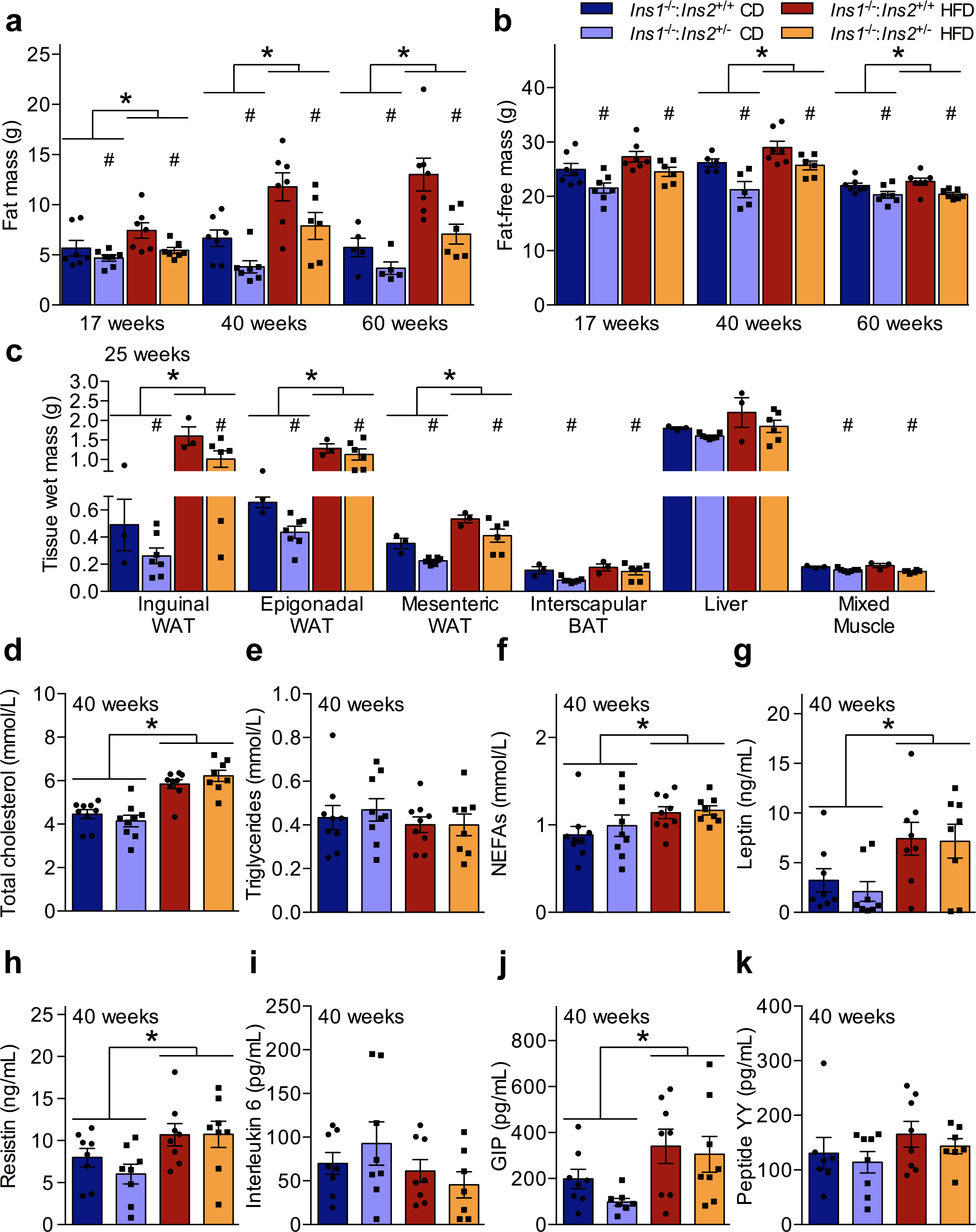
Attenuated Adiposity and Fat-free Mass in Cohort B *Ins1*^−/−^:*Ins2*^+/−^ Males. Periodic DEXA-measured (**a**) fat mass and (**b**) fat-free mass is shown in cohort B male mice (n = 5−7). This corresponded to (**c**) wet masses of inguinal, epigonadal, and mesenteric WAT depots, the interscapular BAT depot, whole liver, and the triceps surae hindlimb mixed muscle group, with all tissues from a group of 25 week-old male mice (n = 3−7) that was separate yet similar in phenotype to cohort B mice. In addition, 4-h fasted (**d**) cholesterol, (**e**) triglycerides, (**f**) non-esterified fatty acids (NEFAs), (**g**) leptin, (**h**) resistin, (**i**) interleukin 6, (**j**) glucose-dependent insulinotropic polypeptide (GIP), and (**k**) peptide YY, is shown in 40 week-old mice from cohort B (n = 7−9). Data are means ± SEM, with scatter points to indicate individual values. Dark blue and red represent CD- and HFD-fed *Ins1*^−/−^:*Ins2*^+/+^ male mice, respectively; pale blue and orange represent CD- and HFD-fed *Ins1*^−/−^:*Ins2*^+/−^ male mice, respectively. *p* ≤ 0.05 denoted by * for CD vs HFD, # for *Ins1*^−/−^:*Ins2*^+/+^ vs *Ins1*^−/−^:*Ins2*^+/−^.

Despite the attenuated adiposity and fat-free mass of cohort B *Ins1*^−/−^:*Ins2*^+/−^ males compared to their *Ins1*^−/−^:*Ins2*^+/+^ littermates, we did not observe notable genotype differences in the fasting levels of circulating lipids and metabolic factors at 40 weeks of age. All HFD-fed mice had higher cholesterol and non-esterified fatty acids than CD-fed animals, without significant differences in triglycerides levels (Fig. 6d-f). In spite of differences in adipose tissue mass, leptin was similarly elevated in all cohort B HFD-fed mice compared to CD-fed mice (Fig. 6g), as was resistin (Fig. 6h). Based on levels of interleukin 6, there did not appear to be significant differences in inflammatory state between groups (Fig. 6i). Glucose-dependent insulinotropic polypeptide levels were similarly elevated in all HFD-fed mice, compared to CD-fed animals (Fig. 6j), but no significant differences were detected in concentrations of peptide YY (Fig. 6k). These data suggest that while a combination of attenuated adiposity and reduced fat-free mass contributed to the smaller body weights of cohort B *Ins1*^−/−^:*Ins2*^+/−^ males compared to their *Ins1*^−/−^:*Ins2*^+/+^ littermate controls, cohort B HFD-fed *Ins1*^−/−^:*Ins2*^+/−^ males nonetheless displayed many of the expected characteristics of high fat feeding.

## Discussion

The initial aim of our work was to test the hypothesis that reducing *Ins2* gene dosage on an *Ins1*-null background would prevent HFD-induced hyperinsulinemia, and thereby protect against obesity in male mice. Contrary to our expectations, inactivating one *Ins2* allele did not cause a consistent reduction of circulating insulin in *Ins1*-null male mice – not even the transient suppression of insulin hypersecretion that was consistently evident in their female *Ins1*^−/−^:*Ins2*^+/−^ littermates at a young age [19]. We report that under some conditions, *Ins1*-null males with reduced *Ins2* mRNA were capable of producing nearly equivalent circulating insulin levels as *Ins1*^−/−^:*Ins2*^+/+^ males, albeit possibly with subtle differences in secretory patterns that could have contributed to modest glucose intolerance. This clearly distinguishes these *Ins1*^−/−^:*Ins2*^+/−^ males from the *Ins2*-null male mice with reduced dosage of the *Ins1* gene, as *Ins1*^+/−^:*Ins2*^+/−^ male mice experienced a sustained suppression of hyperinsulinemia [14]. Our findings show that *Ins1*^−/−^:*Ins2*^+/−^ male mice exhibit phenotypic hyper-variability across cohorts with respect to insulin levels, glucose homeostasis, and weight gain with chronic high fat feeding.

In the current study, all HFD-fed mice in the first experimental cohort, cohort A, showed notable insulin hypersecretion and weight gain, without significant effects of reduced *Ins2* dosage. In contrast, cohort B tended towards a less pronounced degree of HFD-induced insulin hypersecretion and peripheral insulin resistance. In addition, in cohort B there seemed to be a sustained reduction in insulin levels and body mass in *Ins1*^−/−^:*Ins2*^+/−^ mice compared to their *Ins1*^−/−^:*Ins2*^+/+^ littermate controls, without detected changes in food intake or energy expenditure. These two cohorts from the same colony were studied approximately one year apart, under consistent experimental conditions in a controlled SPF facility. Despite this, by one year of age the average differences in fasting insulin levels between the two cohorts were considerably more pronounced than the difference between having one or two functional *Ins2* alleles (in either cohort).

It is important to note that pronounced phenotypic variability between cohorts of *Ins1*^−/−^:*Ins2*^+/−^ male mice, particularly with respect to body mass, was also evident within another animal facility. We cannot explain the widely diverse phenotypic responses to reduced *Ins2* gene dosage in male mice. However, it is clear that cross-cohort phenotypic variability in *Ins1*^−/−^:*Ins2*^+/−^ males has been observed in two distinct facilities to a degree that was not observed in their female littermates [19], nor in *Ins2*-null male or female mice with full or partial *Ins1* expression [14], despite the fact that these similar mouse models were studied by our group in the same time frames and under similar conditions [14,19]. Therefore, in *Ins1*-null male mice, the phenotypic outcomes of *Ins2* gene modulation appear to be susceptible to a wide range in variability.

Phenotypic variability, in general, is poorly understood, but likely affects many long-term animal studies. There is evidence that the *in utero* and neonatal environments (*e.g.* [24–27]), gut microbiome composition (*e.g.* [28–30]), and exposure to different stressors, including temperature (*e.g.* [31–33]), noise (*e.g.* [34,35]), social hierarchy (*e.g.* [36,37]), and even the sex of the researchers working with animal subjects [38], can have far-reaching effects on many physiological parameters. Additional considerations include animal background strain or sub-strain, and genetic drift within a colony (*e.g.* [2,39–42]). These variables can confound experimental results through such means as altering the endocrine milieu, or changing gene expression levels, directly or via the epigenome. However, it should be noted that we attempted to control for many of these potentially confounding variables at a reasonable level in our investigation.

There are numerous other factors that may have played a part in the observed phenotypic heterogeneities in our investigation. For instance, the murine *Ins2* gene has been shown to be subject to developmental stage-dependent and tissue-specific genomic imprinting [11,43,44]. We considered the possibility that the disrupted *Ins2* allele may have had variable effects depending on whether it was inherited from the maternal or the paternal side, particularly as our mice lacked the potential for compensatory *Ins1* expression. Although we did not observe obvious, consistent parental effects on the experimental animals, genomic imprinting is a complex system that has not yet been fully elucidated (see [44–46]), and a potential role cannot be fully ruled out.

As the experimental cohorts were separated temporally, another potential explanation for cross-cohort variability is subtle environmental changes across the years (although the cohorts were roughly matched for seasons, since they were approximately one year apart). There were no differences between cohorts with respect to the average number of siblings sharing their *in utero* and neonatal environments, nor in numerous controlled parameters. However, in long-term experiments it is not always possible to avoid such environmental perturbations as minor earthquakes, construction periods around a facility, and so forth. At 27 weeks, an age when it was becoming clear that the two cohorts were diverging in their patterns of insulin secretion and weight gain, a single blood sample per mouse provided limited indication that levels of the stress hormone corticosterone might have been slightly higher in cohort B males, compared to mice from cohort A. Effects may vary depending on duration or type of stressor, but there is evidence that chronic stress [47,48] or glucocorticoid exposure itself [49–52] can lead to reduced insulin secretion in rodents. Interestingly, the glucocorticoid receptor has been shown to bind to a negative regulatory element of the human *INS* gene [53]. In our results, there was a negative correlation between basal insulin levels and corticosterone across both cohorts in HFD-fed *Ins1*^−/−^:*Ins2*^+/−^ males. Interestingly, an under-powered, rudimentary assessment of plasma samples available from prior experiments showed that 27 week-old HFD-fed *Ins1*^−/−^:*Ins2*^+/+^ and *Ins1*^−/−^:*Ins2*^+/−^ males in a conventional animal facility (see accompanying article) tended to have even higher corticosterone levels and lower fasting insulin levels (data not shown) than in the SPF facility. We suggest that in the current study, reduced exposure to stress, signified by decreased plasma corticosterone, may have partially accounted for some HFD-fed *Ins1*^−/−^:*Ins2*^+/−^ male mice having the ability to produce nearly equivalent amounts of fasting insulin as their *Ins1*^−/−^:*Ins2*^+/+^ littermates. However, multiple other contributing factors likely influenced these outcomes.

The hypothetically environmental-dependent or stress-dependent ability to compensate for reduced *Ins2* dosage at the level of insulin translation and/or secretion was only observed in male *Ins1*^−/−^:*Ins2*^+/−^ mice. Although the female littermates experienced largely consistent conditions under the same experimental time frame, they did not exhibit similar patterns in insulin levels. These findings thereby point to the possibility of sex-specific regulation of murine insulin 2 production or secretion. Both testosterone and estrogen have the capacity to stimulate insulin production and secretion [54–56], so if the gonadal steroids have differential effects on *Ins2* and or insulin 2 peptide, it might have contributed to the observed male-specific variability in insulin levels for *Ins1*^−/−^:*Ins2*^+/−^ mice. However, there are numerous other sex-specific physiological differences that could have also played a role.

Overall, results from this investigation should serve to highlight the range of insulin’s physiological effects, and the phenotypic characteristics that can change in association with variability in insulin levels. We were unable to properly test the hypothesis that prevention of HFD-induced hyperinsulinemia would protect against obesity using this male mouse model, due to non-uniform effects of reducing *Ins2* gene dosage on circulating insulin in *Ins1*-null males. However, we did observe a positive correlation between body mass and fasting insulin, which supports the concept that HFD-fed mice with lower endogenously produced circulating insulin also tend to have reduced obesity [14,19]. In addition, in cohort B experimental mice we show that reduced *Ins2* expression and subtle reductions in insulin levels can be associated with a long-term attenuation of whole body growth, including both fat-free mass and adipose tissue. While it might be expected that differences in insulin levels would primarily affect circulating glucose and other metabolites, we found that anabolic repercussions could be detected in the absence of significant alterations to glucose homeostasis.

Our study shows that circulating insulin levels varied widely in male mice lacking the potentially stabilizing effects of *Ins1* expression. Indeed, measured insulin levels in humans and wildtype mice have been shown to be subject to considerable variation [2–4], and our results suggest that inadvertent stress exposure may be one factor underlying variability in insulin levels. In addition, we show that it is important to look beyond insulin’s well-characterized effects on glucose homeostasis when evaluating the physiological effects of divergent insulin levels.

## Acknowledgements

The authors acknowledge Xiaoke Hu and Subashini Karunakaran for assistance with animal studies. In addition, Xiaoke Hu and Farnaz Taghizadeh provided assistance with the tissue harvests and islet perifusion experiments. We thank Susanne M. Clee for providing use of the metabolic cages, and for critical manuscript input. Timothy J. Kieffer is acknowledged for input on study design and providing use of the Luminex multiplexing instrument.

## References

1. Fu Z, Gilbert ER, Liu D. Regulation of insulin synthesis and secretion and pancreatic Beta-cell dysfunction in diabetes. Curr Diabetes Rev. 2013;9: 25–53.

2. Berglund ED, Li CY, Poffenberger G, Ayala JE, Fueger PT, Willis SE, et al. Glucose metabolism in vivo in four commonly used inbred mouse strains. Diabetes. 2008;57: 1790–1799.

3. McAuley KA, Williams SM, Mann JI, Walker RJ, Lewis-Barned NJ, Temple LA, et al. Diagnosing insulin resistance in the general population. Diabetes Care. 2001;24: 460–464.

4. Li C, Ford ES, McGuire LC, Mokdad AH, Little RR, Reaven GM. Trends in hyperinsulinemia among nondiabetic adults in the U.S. Diabetes Care. 2006;29: 2396–2402.

5. Schumacher MC, Hasstedt SJ, Hunt SC, Williams RR, Elbein SC. Major gene effect for insulin levels in familial NIDDM pedigrees. Diabetes. 1992;41: 416–423.

6. Mayer EJ, Newman B, Austin MA, Zhang D, Quesenberry CP, Edwards K, et al. Genetic and environmental influences on insulin levels and the insulin resistance syndrome: an analysis of women twins. Am J Epidemiol. 1996;143: 323–332.

7. Soares MB, Schon E, Henderson A, Karathanasis SK, Cate R, Zeitlin S, et al. RNA-mediated gene duplication: the rat preproinsulin I gene is a functional retroposon. Mol Cell Biol. 1985;5: 2090–2103.

8. Davies PO, Poirier C, Deltour L, Montagutelli X. Genetic reassignment of the Insulin-1 (*Ins1*) gene to distal mouse chromosome 19. Genomics. 1994;21: 665–667.

9. Wentworth BM, Schaefer IM, Villa-Komaroff L, Chirgwin JM. Characterization of the two nonallelic genes encoding mouse preproinsulin. J Mol Evol. Springer; 1986;23: 305–312.

10. Deltour L, Leduque P, Blume N, Madsen O, Dubois P, Jami J, et al. Differential expression of the two nonallelic proinsulin genes in the developing mouse embryo. Proc Natl Acad Sci USA. 1993;90: 527–531.

11. Deltour L, Montagutelli X, Guenet J-L, Jami J, Páldi A. Tissue- and developmental stage-specific imprinting of the mouse proinsulin gene, Ins2. Dev Biol. 1995;168: 686–688.

12. Hay CW, Docherty K. Comparative analysis of insulin gene promoters: implications for diabetes research. Diabetes. 2006;55: 3201–3213.

13. Meur G, Qian Q, da Silva Xavier G, Pullen TJ, Tsuboi T, McKinnon C, et al. Nucleo-cytosolic shuttling of FoxO1 directly regulates mouse Ins2 but not Ins1 gene expression in pancreatic beta cells (MIN6). J Biol Chem. 2011;286: 13647–13656.

14. Mehran AE, Templeman NM, Brigidi GS, Lim GE, Chu K-Y, Hu X, et al. Hyperinsulinemia drives diet-induced obesity independently of brain insulin production. Cell Metab. 2012;16: 723–737.

15. Linde S, Nielsen JH, Hansen B, Welinder BS. Reversed-phase high-performance liquid chromatographic analyses of insulin biosynthesis in isolated rat and mouse islets. J Chromatogr. 1989;462: 243–254.

16. Wentworth BM, Rhodes C, Schnetzler B, Gross DJ, Halban PA, Villa-Komaroff L. The ratio of mouse insulin I:insulin II does not reflect that of the corresponding preproinsulin mRNAs. Mol Cell Endocrinol. 1992;86: 177–186.

17. Shiao M-S, Liao B-Y, Long M, Yu H-T. Adaptive evolution of the insulin two-gene system in mouse. Genetics. 2008;178: 1683–1691.

18. Leroux L, Desbois P, Lamotte L, Duvilli B, Cordonnier N, Jackerott M, et al. Compensatory responses in mice carrying a null mutation for Ins1 or Ins2. Diabetes. 2001;50: 150S–153.

19. Templeman NM, Clee SM, Johnson JD. Suppression of hyperinsulinaemia in growing female mice provides long-term protection against obesity. Diabetologia. 2015; doi: 10.1007/s00125-015-3676-7

20. Duvillié B, Cordonnier N, Deltour L, Dandoy-Dron F, Itier JM, Monthioux E, et al. Phenotypic alterations in insulin-deficient mutant mice. Proc Natl Acad Sci USA. 1997;94: 5137–5140.

21. Lee KTY, Karunakaran S, Ho MM, Clee SM. PWD/PhJ and WSB/EiJ mice are resistant to diet-induced obesity but have abnormal insulin secretion. Endocrinology. 2011;152: 3005–3017.

22. Salvalaggio PRO, Deng S, Ariyan CE, Millet I, Zawalich WS, Basadonna GP, et al. Islet filtration: a simple and rapid new purification procedure that avoids ficoll and improves islet mass and function. Transplantation. 2002;74: 877–879.

23. Dror V, Nguyen V, Walia P, Kalynyak TB, Hill JA, Johnson JD. Notch signalling suppresses apoptosis in adult human and mouse pancreatic islet cells. Diabetologia. 2007;50: 2504–2515.

24. O’Regan D, Kenyon CJ, Seckl JR, Holmes MC. Glucocorticoid exposure in late gestation in the rat permanently programs gender-specific differences in adult cardiovascular and metabolic physiology. Am J Physiol Endocrinol Metab. 2004;287: E863–70.

25. Fernandez-Twinn DS, Ozanne SE. Mechanisms by which poor early growth programs type-2 diabetes, obesity and the metabolic syndrome. Physiol Behav. 2006;88: 234–243.

26. Murgatroyd C, Patchev AV, Wu Y, Micale V, Bockmühl Y, Fischer D, et al. Dynamic DNA methylation programs persistent adverse effects of early-life stress. Nat Neurosci. 2009;12: 1559–1566.

27. Maniam J, Antoniadis C, Morris MJ. Early-Life Stress, HPA Axis Adaptation, and Mechanisms Contributing to Later Health Outcomes. Front Endocrinol (Lausanne). 2014;5: 73.

28. Backhed F, Ding H, Wang T, Hooper LV, Koh GY, Nagy A, et al. The gut microbiota as an environmental factor that regulates fat storage. Proc Natl Acad Sci USA. 2004;101: 15718–15723.

29. Cani PD, Bibiloni R, Knauf C, Waget A, Neyrinck AM, Delzenne NM, et al. Changes in gut microbiota control metabolic endotoxemia-induced inflammation in high-fat diet-induced obesity and diabetes in mice. Diabetes. 2008;57: 1470–1481.

30. Bruce-Keller AJ, Salbaum JM, Luo M, Blanchard E, Taylor CM, Welsh DA, et al. Obese-type Gut Microbiota Induce Neurobehavioral Changes in the Absence of Obesity. Biol Psychiatry. 2015;77: 607–615.

31. Thurlby PL, Trayhurn P. The role of thermoregulatory thermogenesis in the development of obesity in genetically-obese (ob/ob) mice pair-fed with lean siblings. Br J Nutr. 1979;42: 377–385.

32. Jhaveri KA, Trammell RA, Toth LA. Effect of environmental temperature on sleep, locomotor activity, core body temperature and immune responses of C57BL/6J mice. Brain Behav Immun. 2007;21: 975–987.

33. Feldmann HM, Golozoubova V, Cannon B, Nedergaard J. UCP1 ablation induces obesity and abolishes diet-induced thermogenesis in mice exempt from thermal stress by living at thermoneutrality. Cell Metab. 2009;9: 203–209.

34. Rasmussen S, Glickman G, Norinsky R, Quimby FW, Tolwani RJ. Construction noise decreases reproductive efficiency in mice. J Am Assoc Lab Anim Sci. 2009;48: 363–370.

35. Pascuan CG, Uran SL, Gonzalez-Murano MR, Wald MR, Guelman LR, Genaro AM. Immune alterations induced by chronic noise exposure: comparison with restraint stress in BALB/c and C57Bl/6 mice. J Immunotoxicol. 2014;11: 78–83.

36. Ebbesen P, Villadsen JA, Villadsen HD, Heller KE. Effect of social order, lack of social hierarchy, and restricted feeding on murine survival and virus leukemia. Ann N Y Acad Sci. 1992;673: 46–52.

37. Sterlemann V, Ganea K, Liebl C, Harbich D, Alam S, Holsboer F, et al. Long-term behavioral and neuroendocrine alterations following chronic social stress in mice: implications for stress-related disorders. Horm Behav. 2008;53: 386–394.

38. Sorge RE, Martin LJ, Isbester KA, Sotocinal SG, Rosen S, Tuttle AH, et al. Olfactory exposure to males, including men, causes stress and related analgesia in rodents. Nat Methods. 2014;11: 629–632.

39. Phillips TJ, Hen R, Crabbe JC. Complications associated with genetic background effects in research using knockout mice. Psychopharmacology (Berl). 1999;147: 5–7.

40. Wolfer DP, Crusio WE, Lipp HP. Knockout mice: simple solutions to the problems of genetic background and flanking genes. Trends in Neurosciences. 2002;25: 336–340.

41. Goren HJ, Kulkarni RN, Kahn CR. Glucose homeostasis and tissue transcript content of insulin signaling intermediates in four inbred strains of mice: C57BL/6, C57BLKS/6, DBA/2, and 129X1. Endocrinology. 2004;145: 3307–3323.

42. Mekada K, Abe K, Murakami A, Nakamura S, Nakata H, Moriwaki K, et al. Genetic differences among C57BL/6 substrains. Exp Anim. 2009;58: 141–149.

43. Giddings SJ, King CD, Harman KW, Flood JF, Carnaghi LR. Allele specific inactivation of insulin 1 and 2, in the mouse yolk sac, indicates imprinting. Nat Genet. 1994;6: 310–313.

44. Duvillié B, Bucchini D, Tang T, Jami J, Pāldi A. Imprinting at the Mouse Ins2 Locus: Evidence for cis- and trans-Allelic Interactions. Genomics. Elsevier; 1998;47: 52–57.

45. Peters J. The role of genomic imprinting in biology and disease: an expanding view. Nat Rev Genet. 2014;15: 517–530.

46. Mott R, Yuan W, Kaisaki P, Gan X, Cleak J, Edwards A, et al. The Architecture of Parent-of-Origin Effects in Mice. Cell. 2014;156: 332–342.

47. Zardooz H, Zahedi Asl S, Naseri MG. Effect of chronic psychological stress on insulin release from rat isolated pancreatic islets. Life Sciences. 2006;79: 57–62.

48. Finger BC, Dinan TG, Cryan JF. High-fat diet selectively protects against the effects of chronic social stress in the mouse. Neuroscience. 2011;192: 351–360.

49. Khan A, Ostenson CG, Berggren PO, Efendic S. Glucocorticoid increases glucose cycling and inhibits insulin release in pancreatic islets of ob/ob mice. Am J Physiol. 1992;263: E663–6.

50. Delaunay F, Khan A, Cintra A, Davani B, Ling ZC, Andersson A, et al. Pancreatic beta cells are important targets for the diabetogenic effects of glucocorticoids. J Clin Invest. 1997;100: 2094–2098.

51. Lambillotte C, Gilon P, Henquin JC. Direct glucocorticoid inhibition of insulin secretion. An in vitro study of dexamethasone effects in mouse islets. J Clin Invest. 1997;99: 414–423.

52. Jeong I-K, Oh S-H, Kim B-J, Chung J-H, Min Y-K, Lee M-S, et al. The effects of dexamethasone on insulin release and biosynthesis are dependent on the dose and duration of treatment. Diabetes Res Clin Pract. 2001;51: 163–171.

53. Goodman PA, Medina-Martinez O, Fernandez-Mejia C. Identification of the human insulin negative regulatory element as a negative glucocorticoid response element. Mol Cell Endocrinol. 1996;120: 139–146.

54. Morimoto S, Fernandez-Mejia C, Romero-Navarro G, Morales-Peza N, Díaz-Sánchez V. Testosterone effect on insulin content, messenger ribonucleic acid levels, promoter activity, and secretion in the rat. Endocrinology. 2001;142: 1442–1447.

55. Alonso-Magdalena P, Ropero AB, Carrera MP, Cederroth CR, Baquié M, Gauthier BR, et al. Pancreatic insulin content regulation by the estrogen receptor ER alpha. PLoS ONE. 2008;3: e2069–e2069.

56. Wong WPS, Tiano JP, Liu S, Hewitt SC, Le May C, Dalle S, et al. Extranuclear estrogen receptor-alpha stimulates NeuroD1 binding to the insulin promoter and favors insulin synthesis. Proc Natl Acad Sci USA. 2010;107: 13057–13062.

